# Effects of short-term plasticity in early olfactory information processing in *Drosophila*

**DOI:** 10.1101/2021.05.02.442289

**Authors:** Yuxuan Liu, Qianyi Li, Chao Tang, Shanshan Qin, Yuhai Tu

## Abstract

In *Drosophila*, olfactory information received by the olfactory receptor neurons (ORNs) is first processed by an incoherent feed forward neural circuit in the antennal lobe (AL) that consists of ORNs (input), the inhibitory local neurons (LNs), and projection neurons (PNs). This “early” olfactory information process has two important characteristics. First, response of a PN to its cognate ORN is normalized by the overall activity of other ORNs, a phenomenon termed “divisive normalization”. Second, PNs respond strongly to the onset of ORN activities, but they adapt to prolonged or continuously increasing inputs. Despite the importance of these characteristics for learning and memory, their underlying mechanism remains not fully understood. Here, we develop a circuit model for describing the ORN-LN-PN dynamics by including key features of neuron-neuron interactions, in particular short-term plasticity (STP) and presynaptic inhibition (PI). Our model shows that STP is critical in shaping PN’s steady-state response properties. By fitting our model to experimental data quantitatively, we found that strong and balanced short-term facilitation (STF) and short-term depression (STD) in STP is crucial for the observed nonlinear divisive normalization in *Drosophila*. By comparing our model with the observed adaptive response to time-varying signals quantitatively, we find that both STP and PI contribute to the highly adaptive response with the latter being the dominant factor for a better fit with experimental data. Our model not only helps reveal the mechanisms underlying two main characteristics of the early olfactory process, it can also be used to predict the PN responses to arbitrary time-dependent signals and to infer microscopic properties of the circuit (such as the strengths of STF and STD) from the measured input-output relation.

## Introduction

Sensory systems have evolved different strategies to efficiently represent and process physiologically relevant stimuli in the presence of various biophysical constraints. For example, the olfactory system is confronted with the challenge that there are numerous odors each consisting of multiple volatile molecules with a wide range of concentrations. Yet the olfactory system possesses a remarkable ability to detect and discriminate odors using a relatively small repertoire of odor receptors (ORs) through a combinatorial code, i.e., each odorant is sensed by multiple receptors and each receptor can be activated by many odorants (1–3).

The functional organization of the olfactory systems across different species is highly conserved (4–6). In both insects and vertebrates, an olfactory receptor neuron (ORN) typically expresses only one type of OR. ORNs that express the same OR converge to the same glomerulus in the olfactory bulb (vertebrates) or antennal lobe (AL, insects). In *Drosophila*, peripheral odor information is processed in AL before transmitted to higher brain areas by projection neurons (PNs) (7, 8). Each PN typically innervates one glomerulus. The transfer function between ORN and PN is a saturating nonlinear function (9–11), i.e., a small ORN input is disproportionally amplified while a strong input saturates the response. Lateral inhibition by the local interneurons (LNs) in the AL increases the level of ORN input needed to drive PNs to saturation, the strength of inhibition scales with the total forward input to the AL, a phenomenon called “divisive normalization” (12, 13), which is beneficial for efficient odor coding (10, 14).

Airborne odors are intermittent and have complex spatio-temporal profiles (15, 16). The ability to detect and respond to temporal variation of odors is crucial for successful odor-guided navigation (17–19). This is partially achieved by the response properties of PNs to time-dependent inputs from ORNs. For example, PNs respond transiently to a step-function like ORN input and fall back to low firing rates for the prolonged input, showing highly adaptive response. For more complex time-dependent ORN input, the response of PNs depends on both the ORN firing rate and its rate of change (20–22). These properties are important for detecting and tracking natural odor stimuli. Although a phenomenological linear-nonlinear model was proposed to fit experimental data (20), a mechanistic understanding of how the ORN-PN-LN circuit in AL leads to the adaptive response is still missing.

The aim of this study is to understand both “divisive normalization” and the adaptive response to time-varying stimuli of PNs by modeling the dynamics of the AL neural circuit. Previous studies showed that synapses between ORNs and PNs exhibit strong short-term plasticity (STP) (9, 23), which is a form of fast activity-dependent modulation of synaptic strength (24–28). Furthermore, inhibition by LNs is due to presynaptic inhibition (PI) at the axon terminal of ORNs (10). In the rest of the paper, we first develop a circuit model of the *Drosophila* AL that incorporates both STP and PI. Then, with our analytical results and numerical simulations, we show that STP is essential for the observed highly nonlinear divisive normalization; and both STP and PI determine the adaptive response observed in experiments.

### A circuit model of the peripheral olfactory system with STP

There are around 50 types of ORNs in *Drosophila* melanogaster, each of them expresses one OR. ORNs that express the same OR converge to the same glomerulus in the AL. A given odor typically activates several types of ORNs, hence different glomeruli. Each PN innervates one glomeulus and projects to higher brain areas like mushroom body and lateral horn. Since most LNs in AL are GABAergic, we will only consider inhibitory interneurons. Although LNs have distinct morphologies, innervation patterns, and response dynamics to odors (29–31), for the purpose of this study, we do not differentiate them in our model. Generally speaking, LNs innervate different glomeruli and target the butons of ORN axons, forming presynatpic inhibition (Fig.1a). We consider the simplified neural circuit of ORN-PN-LN in the antennal lobe, which forms an incoherent feedforward loop (IFFL) as shown in Fig.1b.

**Fig. 1.**
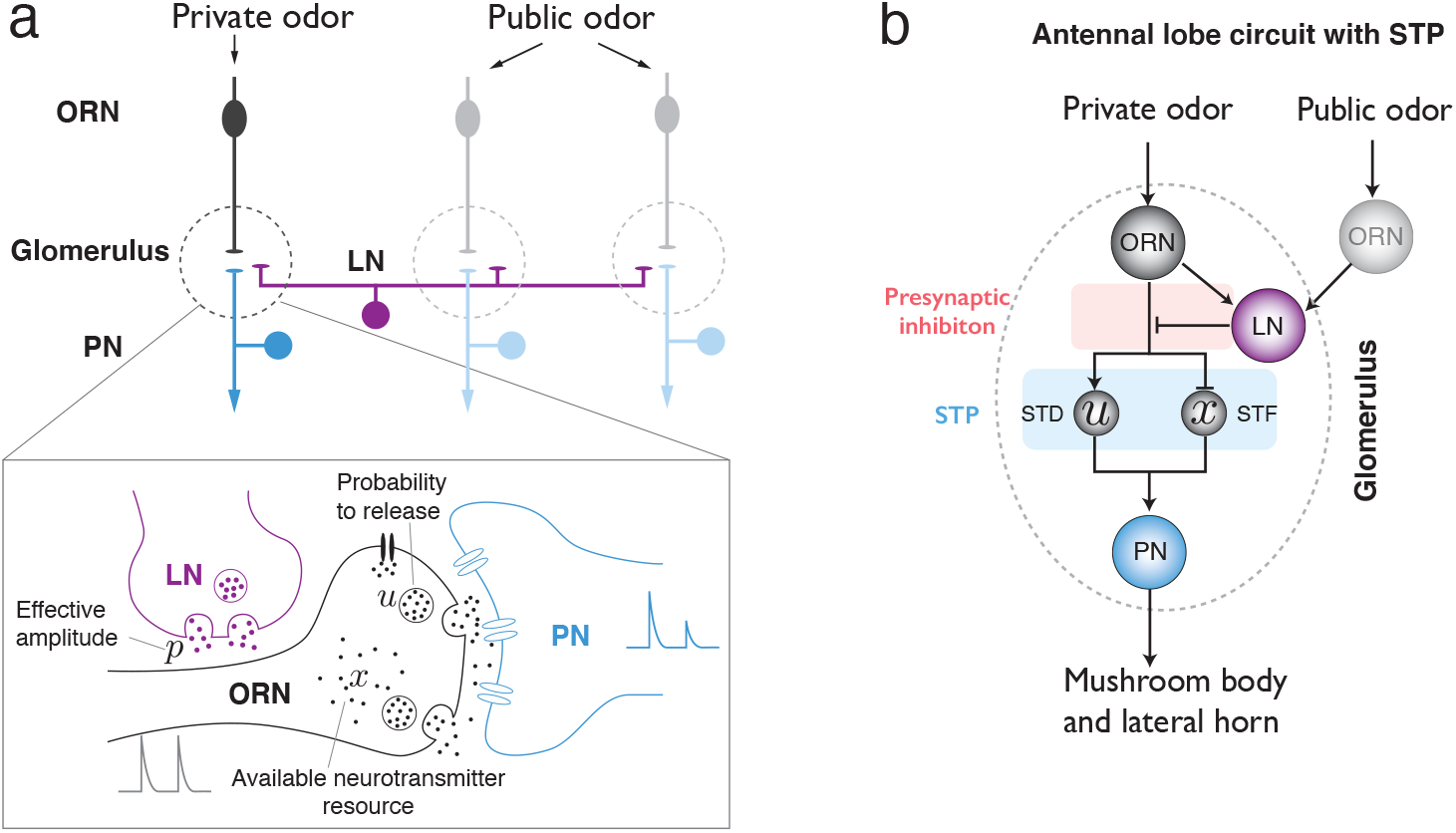
Illustration of the antennal lobe (AL) circuit model. (a) Anatomy of the peripheral olfactory system in *Drosophila*. Olfactory receptor neurons (ORNs) that express the same type olfactory receptors innervate the same glomeurlus (dashed region) in AL. Most projection neurons (PNs) send dendrites into one glomerulus and receive synaptic input from the ORN. Although each glomerulus might be innervated by different PNs, only one PN is shown. Glomeuruli are laterally connected by local neurons (LNs,magenta), which interact with ORNs and PNs. A private odor only activates a specific type of ORNs, while a public odor activates a large number of ORNs that innervate different glomeruli. Lower: close-up of the synaptic interaction between ORN, PN and LN. Both PI and STP are considered in our model. (b) Schematics of the AL circuit with STP effect and PI mediated by LNs.

For simplicity, we assume that the firing rate (*R*_PN(LN)_) of the PN (LN) is proportional to its conductance 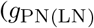 (32): 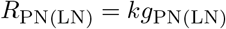 with a constant *k*, which is verified in the leaky integrate-and-fire model (see numerical validation in Fig.S1 in Supplementary Material (SM)). Dynamics of PN (LN) conductance 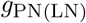 can be written as:

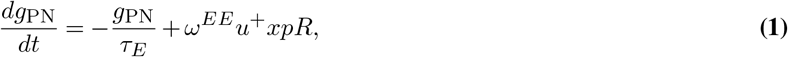

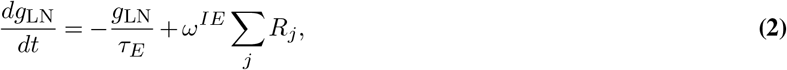

where *R* is the firing rate of the cognate ORN that responds to a particular (private) odorant. The sum Σ_*j*_ in Eq. (2) is over all ORNs connected to the LN, including non-cognate ORNs that respond only to public odorant(s). The timescale *τ_E_* is the relaxation time of conductance. *ω^EE^* and *ω^IE^* are synaptic weights of the synapses from ORN to PN and LN respectively, which we assume to be homogeneous among different ORNs.

The effect of PI is modeled by Eq. (1) with a (dimensionless) variable 0 < *p* < 1 that represents reduction of the effective ORN firing rate due to presynaptic inhibition by LN. The dynamics of *p*, simplified from previous studies (33, 34), is modeled as:

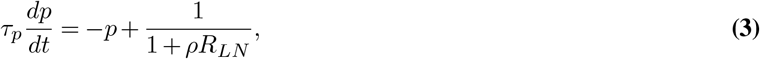

where *ρ* is a constant and *τ_p_* is the relaxation time of *p*. In the limit *τ_p_*≪*τ_E_*, we can use the quasi-steady state approximation 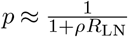, which suggests that *p* decreases with *R*_LN_.

The effect of STP can be separated into short term facilitation (STF) and short term depression (STD), which are modeled by *u*^+^ and *x* in Eq. (1) respectively. Following from previous work (35), we denote *u*^−^ (*u*^+^) as the releasing probability just before (after) the arrival of a presynaptic spike; and *x* as the fraction of available neurotransmitters (Fig. 1a). Applying the mean-field model for STP (35), we have the following dynamics for *u*^±^ and *x*:

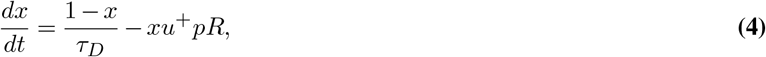

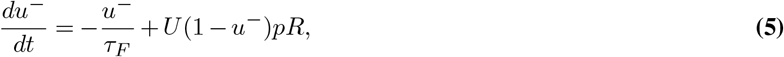

where *u*^+^ = *u*^−^ + *U* (1 − *u*^−^) with *U* as the increment in release probability after each spike. Without any presynaptic firing (*R* = 0), we have *x* = 1, *u*^−^ = 0, and *u*^+^ = *U* at steady state. With presynaptic firing, *x* decreases and *u*^+^ increases before their steady state values are recovered with time constants *τ_D_* and *τ_F_* respectively. Therefore, the strength of STD and STF can be measured by their normalized recovery time (dimensionless), 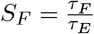 and 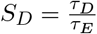. The longer the recovery time, the stronger the effect.

## Results

We use the neural circuit model proposed in the previous section to describe and explain several response properties of PNs including divisive normalization for constant (steady state) inputs and adaptive response to time-varying inputs. In both cases, we compare our model results with existing experiments and focus on understanding the effects of STP and PI on the observed behaviors.

### STP is crucial for the observed nonlinear divisive normalization

For any input (cognate ORN with firing rate *R*) to the peripheral olfactory system, its output, i.e., the firing rate of the cognate PN can be computed by solving Eqs. (1)–(5). For a constant input (fixed firing rates of all ORNs), the steady state PN response can be determined analytically:

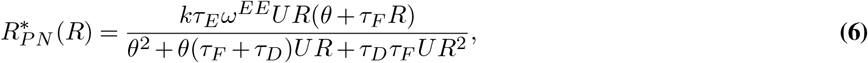

where *θ* ≡ 1 + *A*Σ_*j*_ *R*_*j*_ with *A* ≡ *kρω^IE^τ_E_*, and 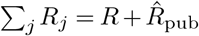 with 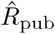 denoting the total input from public ORNs.

In the absence of STP, i.e., when *τ_D_* → 0 and *τ_F_* → 0, *x* = 1 and *u*^+^ = *U* remain constant, the PN response Eq. (6) reduces to:

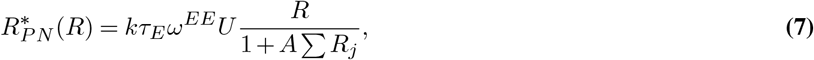

where both the numerator and denominator depend linearly on *R*. Eq. (7) can thus be called “linear” divisive normalization. In the presence of STP, i.e., when *τ_F_* ≠ 0 and *τ_D_* ≠ 0, both the denominator and the numerator in Eq. (6) are nonlinear in *R*, which we call “nonlinear” divisive normalization.

The PN response curve exhibits a sigmoidal shape that can be characterized by three parameters: the maximum response 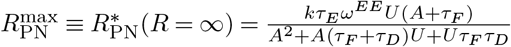; the half maximum input *R* defined as 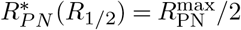; and an effective Hill coefficient 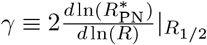. From Eq. (7), we have *γ* = 1 in the absence of STP. In the presence of STP, we have *γ* > 1, which can be used to characterize the gain of the response. In Fig. 2a&b, the PN response function Eq. (6) with different STP strengths are shown. As expected, for a given input *R*, STF enhances the response while STD suppresses it with the latter having a stronger effect. The dependence of the gain *γ* on the STP strengths *S_D_* and *S_F_* are shown in Fig. 2c. Interestingly, the gain parameter *γ* is enhanced by both STD and STF. Larger values of *γ* are reached by having roughly the same STD and STF strengths (dotted line in Fig. 2c).

**Fig. 2.**
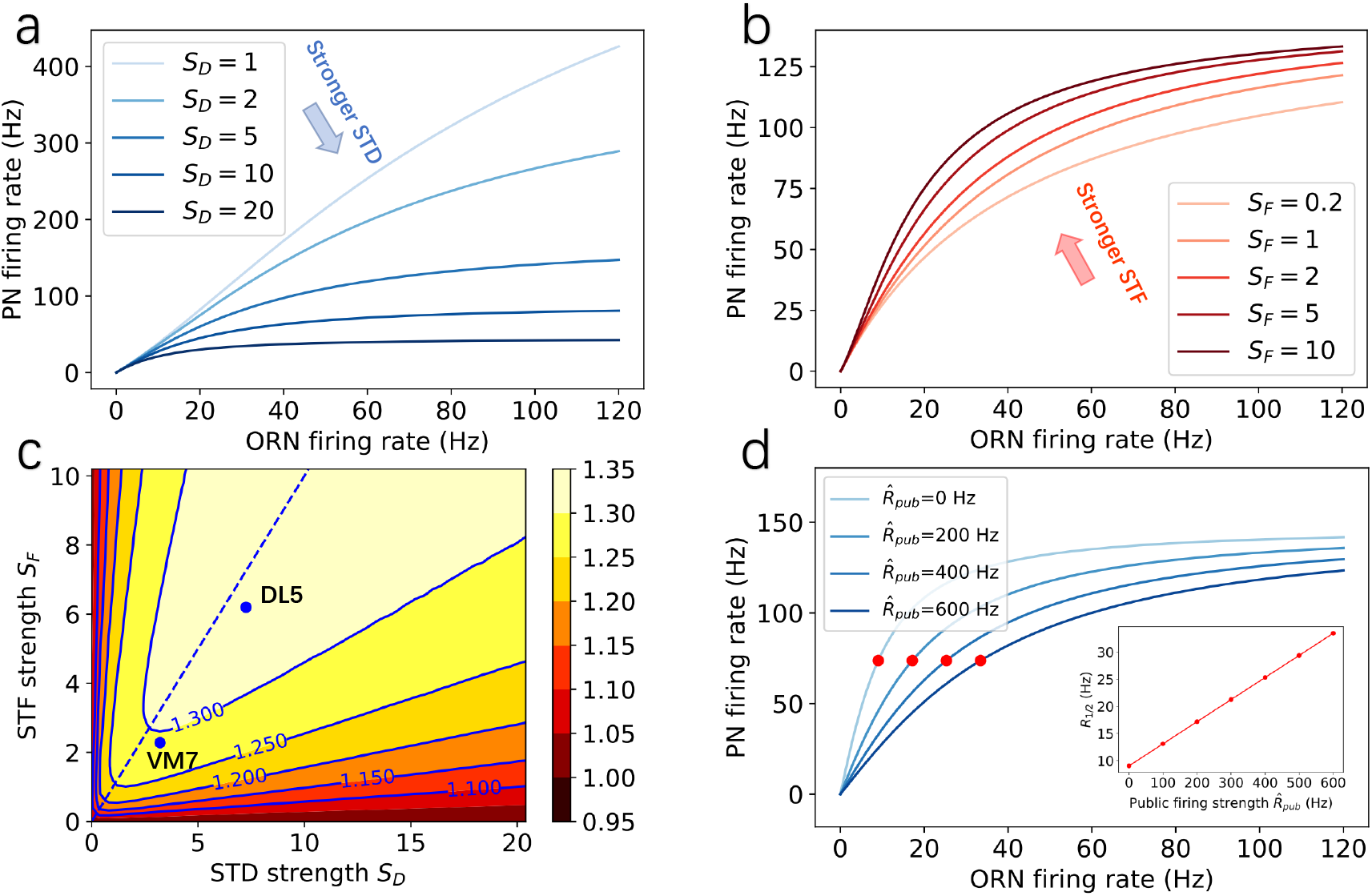
The effects of STP (STF and STD) on the nonlinear divisive normalization effect in PNs’ responses. (a,b) Steady state responses of PN to different cognate ORN firing rates for (a) different STD strengths *S_D_* and (b) different STF strengths *S_F_*. In (a) *S_F_* = 2, in (b) *S_D_* = 6. (c) The dependence of the effective Hill coefficient *γ* on the STF strength (*S_F_*) and the STD strength (*S_D_*). The values of *S_D_* and *S_F_* used in fitting the experimental data for DL5 and VM7 (see Fig. 3) are also shown in the figure. The dotted line corresponds to perfectly balanced STF and STD strength: *S_D_ = S_F_*. (d) Steady state responses of PN to cognate ORN firing rates in the presence of different public firing rates 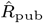. Red dots mark ORN firing rates at the half maximum firing of PNs (*R*_1*/*2_). The inset shows that *R*_1*/*2_ increases linearly with 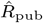. Other model parameters used here are: *U* = 0.24, 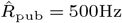, k = 5Hz/nS, *ω*^EE^ = 180nS, *ω*^IE^ = 10nS, *τ*_E_ = 50ms, *ρ* = 0.0018.

As shown in Eq. (6), the PN response is suppressed by the firing of LN, which can be activated by many ORNs besides the cognate ORN. This introduces lateral inhibition and reduces the response to a private odorant in the presence of public odorants (Fig.2d). This divisive normalization effect was fitted by a Hill function in Olsen *et al* (12):

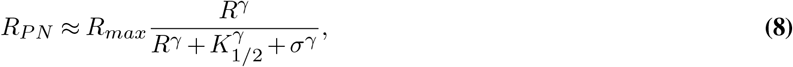

where *σ* is assumed to be proportional to the sum of the firing rates of non-cognate ORNs; *R*_max_ is the maximum PN firing rate; *k*_1*/*2_ is the firing rate of ORN at which PN has the half maximum response when *σ* = 0, *γ* is the Hill coefficient. In (12), the authors were able to fit their experimental data with *γ ≈* 1.5.

From our model, this experimental observation value of *γ*(> 1) suggests the existence of strong STP effects. To verify this hypothesis, we fit the experimental results reported in (12) using Eq. (6) from our model. In a typical experiment, PN responses to a private odorant (which activates only the cognate ORN) with a background of different concentrations of a public odorant (presumably activates many other ORNs) was measured. As shown in Fig. 3, each color represents a different concentration of the public odorant, the exact firing rate is not available from the experiment, but was assumed to be proportional to the measured local field potential (LFP). In our model this is fitted by adjusting the value of total firing rate of public ORNs 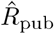.

**Fig. 3.**
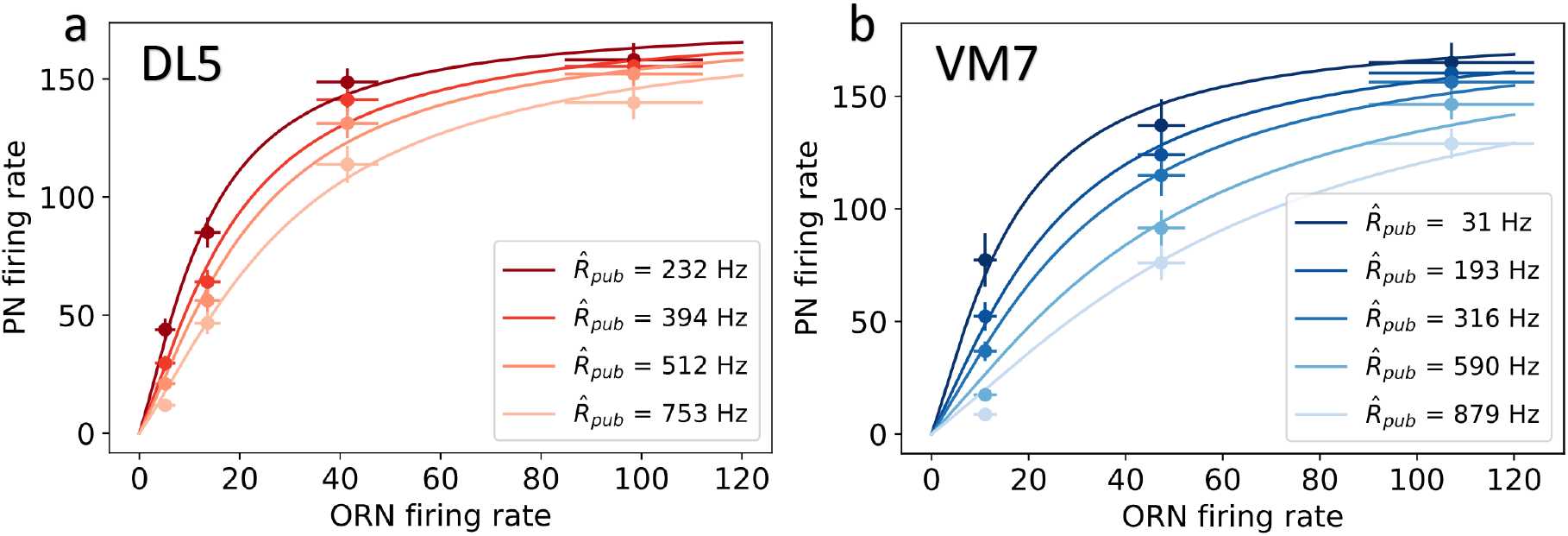
The circuit model with STP and PI can fit experimental data exhibiting the nonlinear divisive normalization effect in a PN’s response. Different shades of colors indicate different strengths of lateral inhibition due to different public ORN firing rates. PNs that inervate the (a) DL5 and (b) VM7 glomeruli are compared. Experimental data are from (12). The best-fit parameters shown in Table 1 are within physiological ranges.

Our model fits well with the measured responses of the PNs that innervate either DL5 or VM7 glomeruli for different levels of public odorant concentrations. The best-fit parameter sets for DL5 PN and VM7 PN are given in Table 1. While most of the parameters for DL5 and VM7 remain approximately the same, there are some differences that are informative. In particular, the two STP timescales (*τ_D_* and *τ_F_*) and the lateral inhibition strength *ρ* are intrinsic properties of the glomerulus and are thus expected to be different. Our model results indicate that both DL5 and VM7 have strong STP effects. Quantitatively, both STD and STF strength are stronger in DL5 than those in VM7, however, the relative strengths of STD and STF, *r* ≡ *τ_D_*/*τ_F_*, remain roughly the same for DL5 (*r* ≈1.1) and VM7 (*r* ≈1.4), which indicates that a strong and balanced STD and STF is responsible for the observed nonlinear divisive normalization in both VM7 and DL5. Both timescales (*τ_D_* and *τ_F_*) obtained from our model fitting are consistent with the range of these timescales measured in experiments (36). We also find that lateral inhibition is slightly stronger in DL5 than that in VM7.

**Table 1.** Parameters in our model used for Fig. 3

| Parameter | Meaning | DL5 | VM7 |
| --- | --- | --- | --- |
| k | linear coefficient from conductance to firing rate | 5 Hz/nS | 5 Hz/nS |
| $\tau_E$ | time constant for excitatory synapse | 50 ms | 50 ms |
| $\omega^{EE}$ | synaptic weight from ORN to PN | 247 nS | 121 nS |
| $\omega^{IE}$ | synaptic weight from ORN to LN | 10 nS | 10 nS |
| $\rho$ | intrinsic strength for presynaptic inhibition | 0.02 | 0.015 |
| U | increase in release probability for facilitation | 0.24 | 0.24 |
| $\tau_D$ | time constant for STD | 345 ms | 161 ms |
| $\tau_F$ | time constant for STF | 309 ms | 114 ms |

### Adaptive response to time-varying signals and its circuit origin

Odors in the environment are highly intermittent and dynamic (15, 16). The ability to detect and respond to temporal variation of odor stimuli is crucial for the survival of many animals. Kim *et al* (20) studied the responses of PNs to different time-dependent ORN signals that follow a triangle-shaped temporal pattern with different peak times. The responses of PNs were “asymmetric”, with a faster rising phase, followed by a plateau and a slower decaying phase, depending on the rate of change in ORN’s firing rate profile (see Fig. 3 in (20)). Here, we use our model to explain the response patterns to these time-dependent signals.

We start by considering the response of PNs to a signal that increases linearly with time, i.e., *R*(*t*) = *Kt*. In the short time limit *t* ≪ *τ_p_*, PI is negligible so the reduction factor *p* ≈ 1 (see Eq. (3)). As a result, the response is linearly proportional to the input 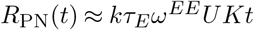. For longer time *t* ≫ *τ_p_*, *p* ≈ (*AR*(*t*))−1 decreases inversely proportional to the input *R*(*t*) which reduces the effective input *pR*(*t*). In fact, the effective input can be approximated as (see SM for detailed derivation):

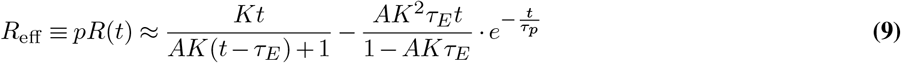

In the long time limit *t* ≫ *τ_E_*, *τ_p_*, the system adapts by adjusting the inhibition factor *p* so that the effective input reaches a constant 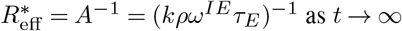 as *t*, which is independent of the input. The corresponding adapted response can thus be determined analytically:

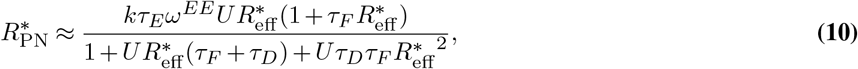

which takes exactly the same form as the response to an effective time-independent signal 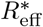 as described in Eq. (6). This is supported by numerical simulation as shown in Fig. 4. The effect of PI is crucial for canceling out the increasing signal, which results to a constant effective input 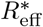. From Eq. (10), it is clear that STP also affects the adapted response 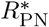 in the same way as it affects the response to a constant signal, i.e., STF enhances the adapted response and STD suppresses it.

**Fig. 4.**
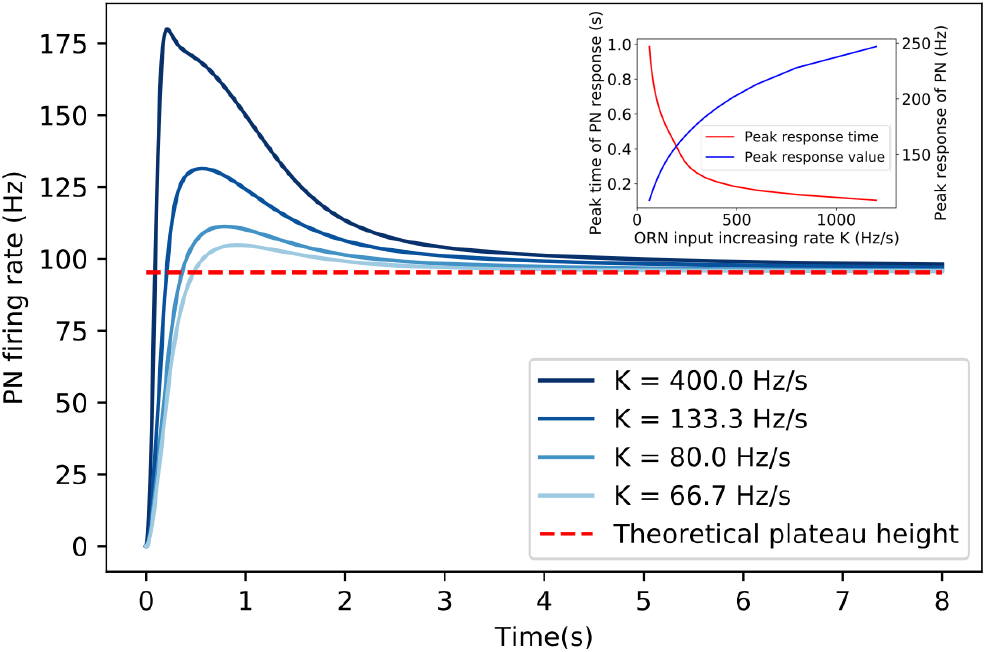
The responses of PN to linearly increasing inputs (ORN firing rate) with different increasing rate *K*. The adapted responses are independent of the increasing rate (*K*) of the input. Dashed line marks the asymptotic response when *t* → ∞ as predicted by Eq. (10). The inset shows how the peak time and peak value of PN response depend on the increasing rate (*K*).

We now study the PN responses to triangle-shaped input signals similar to those used in experiments (20) with our model. As shown in Fig. 5a, the general response dynamics follow closely the experimentally observed behaviors. During the rising phase of the input signal, the PN response activity reaches its peak in a timescale that depends on the rate of change (*K*) of the input signal. For small *K* shown as the purple line in Fig. 5a, the PN response reaches a plateau before the input reaches its peak due to the adaptive effect of presynaptic inhibition described above. The shaded region in Fig. 5a (lower panel) shows the range of the experimentally observed plateau consistent with the model result (purple line). Note that for higher *K* the plateau activities depend on the rates of change in the input signals because they do not have enough time to reach the adapted value. In the descending phase of the input signal, the PN activity decreases following the input.

**Fig. 5.**
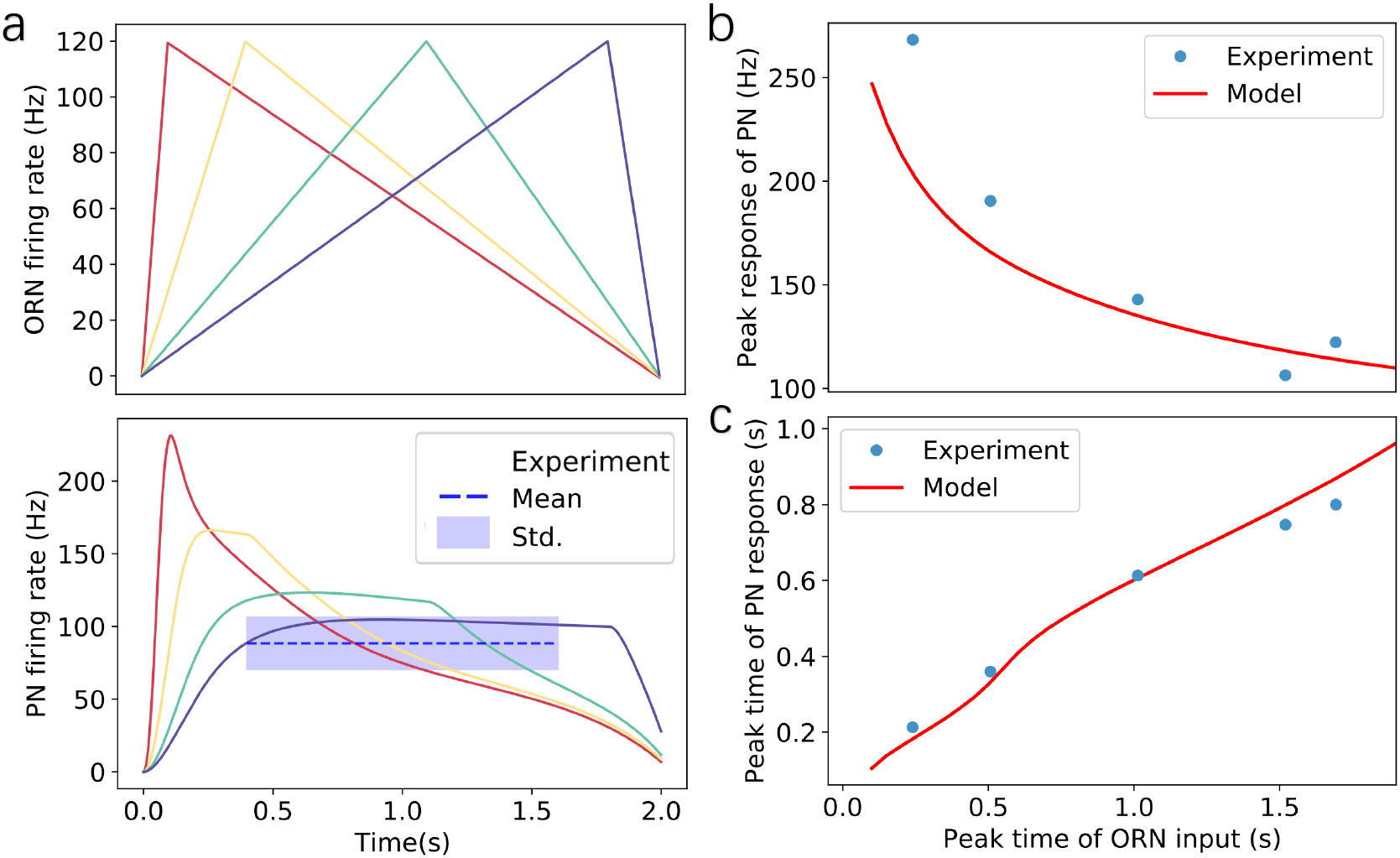
Adaptive responses of PN to triangle-shaped inputs (ORN spike rates). (a) Upper panel: Simulated triangle-shaped ORNs firing rates with different increasing rates in the rising phase. The peak inputs (ORN firing rates) are the same for all cases. Lower panel: responses of PNs to triangle-shaped ORNs inputs. For slow and medium increasing inputs, PNs reach plateau responses. The dotted line shows the average plateau response of PN estimated from experimental data with the shaded region indicating the standard derivation. (b) The peak responses of PNs increase with the rates of change in input signals. Our model result (solid line) agrees well with the experiment (dots). (c) PN’s response reaches a peak earlier than that of the input signal. The model result (solid line) has an excellent agreement with the experiment (dots). Experimental results are from (20). Parameters used in the model: *U* =0.24, *R*_pub_ = 0Hz, k = 5Hz/nS, *ω*^EE^ = 75nS, *ω*^IE^ = 21nS, *τ*_E_ = 55ms, *ρ* = 0.008, *τ*_p_ = 300ms, *τ*_F_ = 50ms, *τ*_D_ = 100ms.

Quantitatively, the PN response can be described by two parameters: the peak response, which is defined as the PN activity at the peak time of the input signal, and the time to reach the peak response. In Fig. 5b&c, we show that the results for these quantities measured from our model are in excellent agreement with those obtained from experiments (20).

In principle, STP (specifically STD) alone can also lead to adaptive responses. However, we find that the adaptive response dynamics to time-varying signals observed in experiments (20) is more strongly affected by PI. This can be seen by comparing PN’s response to triangle-shaped input in four model variants: the standard model used in Fig. 5 with strong PI and moderate STP, a model with PI only without STP (*S_D_* = *S_F_* = 0), and a model with only STP without PI (*ρ* = 0). As shown in Fig. S2 in SM, the model response agrees with experiments qualitatively well even with PI alone, however, STP is needed to achieve quantitative agreement with experiments. On the other hand, even though a very strong STD may also cause the system to adapt (Fig. S2b), the fit of the STP-only model to experiments remains poor even by fine tuning the STF and STD strengths (see Fig. S2b–d in SM for details).

## Discussion

In this study, we developed a simple neural circuit model for the antennal lobe of *Drosophila* and systematically studied the role of STP and PI in early odor information processing. Combining analytical derivations of a steady-state solution of the model and numerical simulations, we showed that the model can capture key characteristics of PNs’ responses to different ORN inputs, in particular divisive normalization and adaptive responses to time-varying signals. Comparison with experimental results revealed that STP is crucial for the observed nonlinear divisive normalization, and the adaptive response to the time-varying ORN input is affected by both STP and PI with the latter playing a more dominant role in the experimental systems we analyzed.

Previous experiments have suggested that STD largely determines the nonlinear response function of PNs (9), and PI plays an important role in divisive normalization (10, 12). Our model extends these studies and showed that STP is crucial for the observed nonlinear divisive normalization. For stable ORN input, STF enhances PN’s response while STD suppresses it. Yet, both STF and STD enhance the nonlinearity effect of the divisive normalization (Fig. 2). Although the transient response of PNs to step-like ORN input has been attributed to STD, our model shows that presynaptic inhibition predominately determines PN’s response to more dynamic ORN input, such as triangle-shaped input patterns (Fig.5). In fact, our model can predict the response properties of PNs to arbitrary time-dependent ORN inputs. For example, when applying a set of sine-wave ORN inputs with different frequencies, our model predicts that the amplitude of PN’s oscillatory response will increase as the frequency gets higher, while the time advance of PN’s peak to ORN’s peak will decrease (see Fig. S3 in SM). These predictions can be tested by future experiments.

Our model not only revealed the underlying mechanisms for the observed nonlinear divisive normalization behavior and adaptive responses to time-varying signals, it also provides a general framework for relating microscopic properties of the system such as time scales and strengths of STP and PI to macroscopic behaviors such as the input-output relation. As demonstrated in this work in the cases of VM7 and DL5 glomeruli, we can use our model to infer/estimate microscopic properties of the system from the measured input-output relation. More specifically, from our model study, the timescales of both STD (*τ_D_*) and STF (*τ_F_*) are predicted to be longer in DL5 than those in VM7, which can be verified by experiments (36). The model-based analysis of the input-output response can be extended to other glomeruli. Our model can also be used to make predictions for changes in the input-output relation when certain microscopic properties, e.g., the STP strength (*τ_F_* and *τ_D_*) or the PI strength (*ρ*) are perturbed. These predictions can be tested in future experiments.

As we focused on building a minimal model to understand the underlying mechanism for nonlinear divisive normalization and adaptive response, we have made several simplifications in our study. At the synapse level, our model ignores the STP effect at the ORN-LN synapses (31), which can affect the LN response. In our circuit model, we only considered one type of LNs for simplicity. There are several type of LNs in the antennal lobe with diverse innervation patterns and physiological properties (29–31). For example, a small fraction of LNs are excitatory. Panglomerular LNs innervate all glomeruli with higher spontaneous firing rates than other LNs, they are inhibited or only weakly excited by odors. Such inhibition of panglomerular LNs tends to dis-inhibit the entire antannal lobe in the presence of odors (29). LNs also show distinct response dynamics to odors. Future study should address how these additional complexity and LN-LN interactions affect PN’s response properties (37). Furthermore, although most PNs innervate a single glomerulus (uPNs) as we studied in this paper, some do receive input from multiple glomeruli (mPNs). uPNs and mPNs have different projection patterns and may carry different aspects of odor information to the higher brain regions such as lateral horn and mushroom body (38). Future studies that incorporate these important features (both at the synapse level and the network level) will further our understanding of the rich dynamics in the early olfactory information processing in *Drosophila*.

## ACKNOWLEDGEMENTS

We thank Guangwei Si, Tianyi Wu and members of Tang lab for helpful discussion. The work by YT is supported by a NIH grant(R35GM131734).

## Supplementary Material

## A. The linear relation between firing rate and conductance

In the main text, we have used the approximation that a neuron’s firing rate is proportional to its conductance 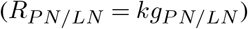 at steady state. Here, we numerically verify the validity of this approximation. Consider a leaky integrate-and-fire neuron whose membrane potential evolves as

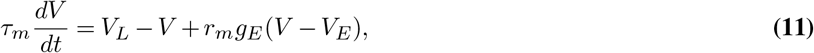

for *V* < *V_th_*. Here *V_L_* and *V_E_* are the resting potential and the reverse potential respectively, *r_m_* is the membrane resistance, *g_E_* is the conductance, *τ_m_* is the membrane time constant and *V_th_* is the firing threshold. When *V* reaches *V_th_*, the membrane potential is reset to *V_L_*. Initializing *V* at the resting potential, i.e., *V* (*t* = 0) = *V_L_*, the solution of Eq. (11) is given by

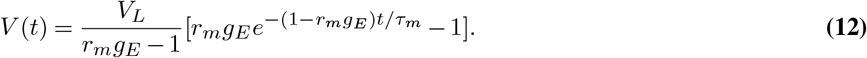

We are interested in the time required Δ*t* for *V* (*t*) to reach the firing threshold *V_th_* from the resting potential *V_L_*. Assuming that *t* ≤ Δ*t* ≪ *τ_m_*, we can expand Eq. (12) linearly in *t*/*τ_m_*:

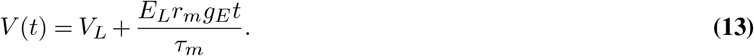

Δ*t* can be solved from above equation:

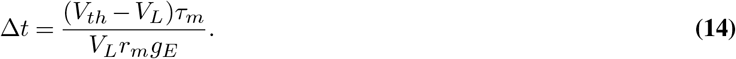

The firing rate of the neuron is defined by how many times *V* reaches *V_th_* in a unit time

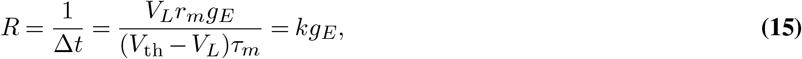

which is proportional to the conductance 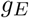. The above approximation is further verified by numerical simulation (Fig. S1).

## B. Response of PNs to triangular-shaped firing rates of ORNs.

In this section, we describe in detail the approximation used in the calculations when we study PN’s response to triangular-shaped inputs. First, we consider a simpler scenario, where ORN’s firing rate increases linearly without bound, i.e., *R = Kt*. Then, Eq.(1–5) in the main text become

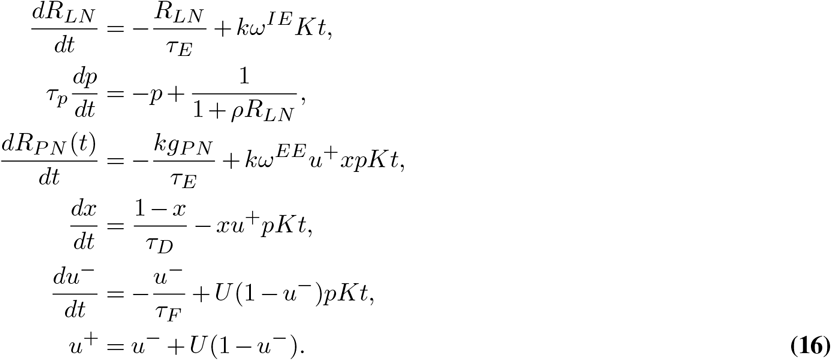

To analyze the adaptive behavior of PN responses, we try to give an approximate solution of the above equations. These equations can be separated into two groups based on the biological mechanisms they describe: the first two equations describe presynaptic inhibition and the rest equations are related to STP.

***The presynaptic inhibition terms.*** Solving the first two equations of Eq. (16) we obtain:

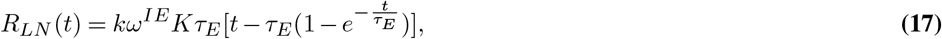

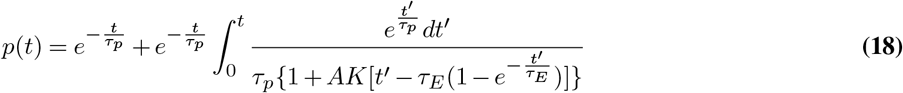

with initial conditions *p*(0) = 1 and *R_LN_* (0) = 0. Here 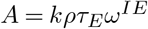, as defined previously in the steady state solution in the main text. Since the plateau appears at *t* ≫ *τ_E_*, the integrand of Eq. (18) can be approximated as

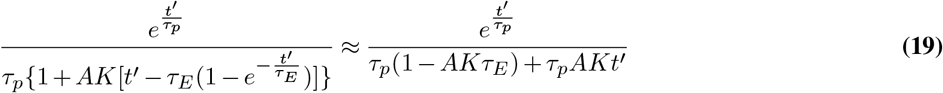

Plug it back into Eq. (18), we obtain:

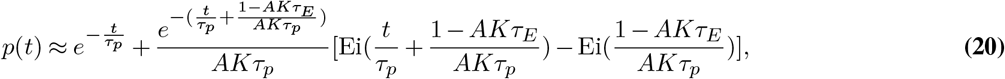

where 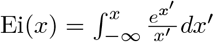 denotes the exponential integral. Asymptotically expand Ei(*x*) as 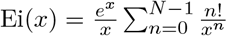 and consider the limit t→∞, we have

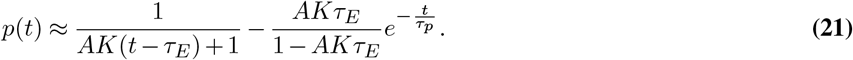

Multiply Eq. (21) by *Kt* gives the effective input *R*_eff_ Eq. (9) in main text.

***The STP terms.*** In this section we consider the last 4 equations in Eq. (16) which describe the STP mechanism. As we discussed in the main text, for *t* ≫ *τ_p_*, *p* ≈ (*AR*(*t*))−1, which leaves the effective input *R*_eff_ = *p*(*t*)*R*(*t*) approximately constant in time. We use this approximation to investigate the plateau behavior of PN response. Plugging it back into Eq. (16), we have 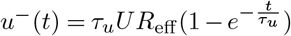 where 1*/τ_u_ ≡*1*/τ_F_* + *R*_eff_ *U*.

The equation for *x* becomes:

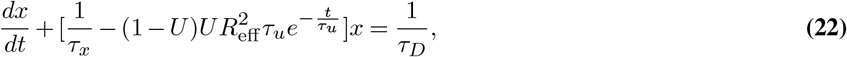

where 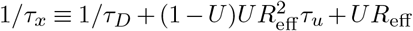. Introducing 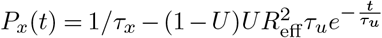,we have

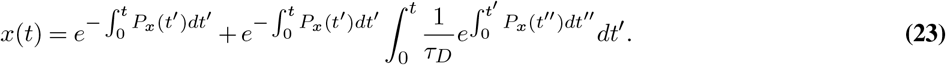

Since we are interested in the plateau behavior of PNs, where *t* ≫ *τ_u_*,*τ_x_*. We can approximate 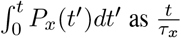. Plug it back to Eq. (23) and neglect higher order corrections, we have *x*(*t*) ≈ *τ_x_/τ_D_*. Similarly,

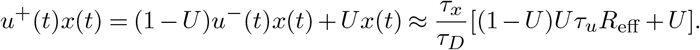

The magnitude of the plateau response of PN can be approximated by its steady state activity for large *t*,

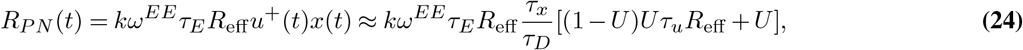

where *R*_eff_ = *p*(*t*)*Kt* ≈ 1/*A* with *p*(*t*) given by Eq. (21). We see that the plateau magnitude of *R_P N_* is related to *R*(*t*) only through *R*_eff_, which is the final value that *pR*(*t*) will reach. It is independent of *t* and *K*, therefore the magnitude of PN response plateau is not affected by the input changing rate *K*, but only depends on properties that are intrinsic to the network.

## C. Model variants incorporating different mechanisms revealing the role of STP in response to time-varying signals.

In this section, we compare several model variant incorporating only part of the mechanisms compared to the full model (standard model) in the main text. As shown in Fig. S2, PN response with only presynaptic inhibition is slightly different from the standard model with both STP and presynaptic inhibition, while the STP only model deviates from the standard model more significantly. In addition, as shown in Fig. S2(b), the STP-only model can not distinguish the experimentally observed different plateau heights for different inputs. We conclude that presynaptic inhibition plays a dominant role in shaping a PN’s response to time varying signals while STP fine tunes the adaptive response.

---

**Fig. S1.**
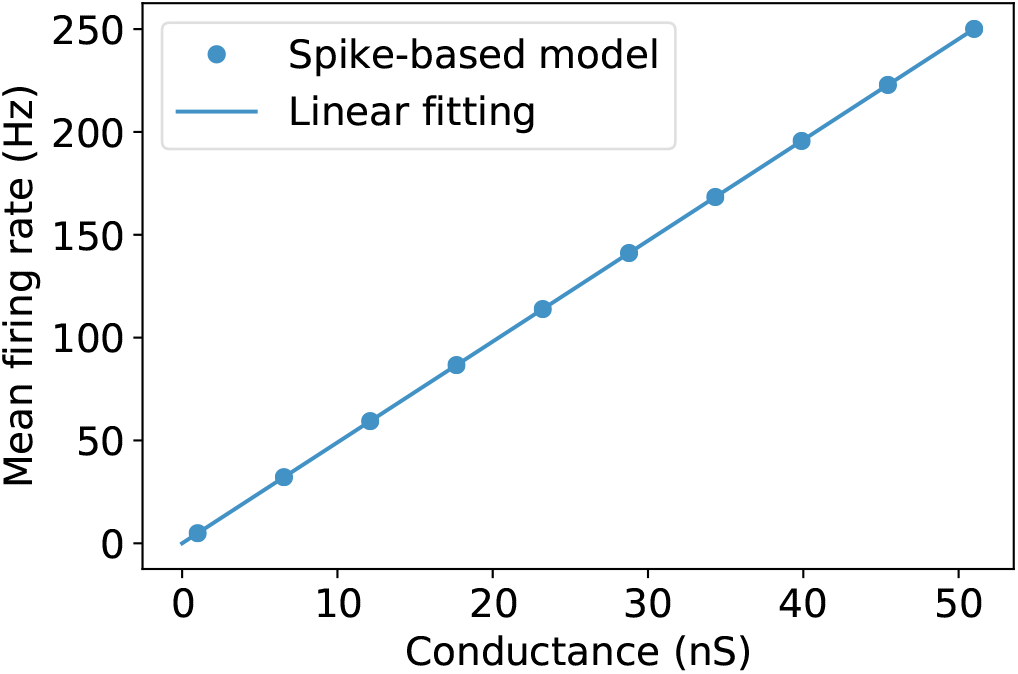
Firing rate is proportional to the steady state conductance (*gE*) in the leaky integrate-and-fire model. The parameters in Eq.11 used for the simulation are: *τ_m_* = 20*ms*, *V_L_* = −70*mV*, *V_E_* = 0*mV*, *V_th_* = −50*mV*, *r_m_* = 3.3 × 10^7^Ω.

**Fig. S2.**
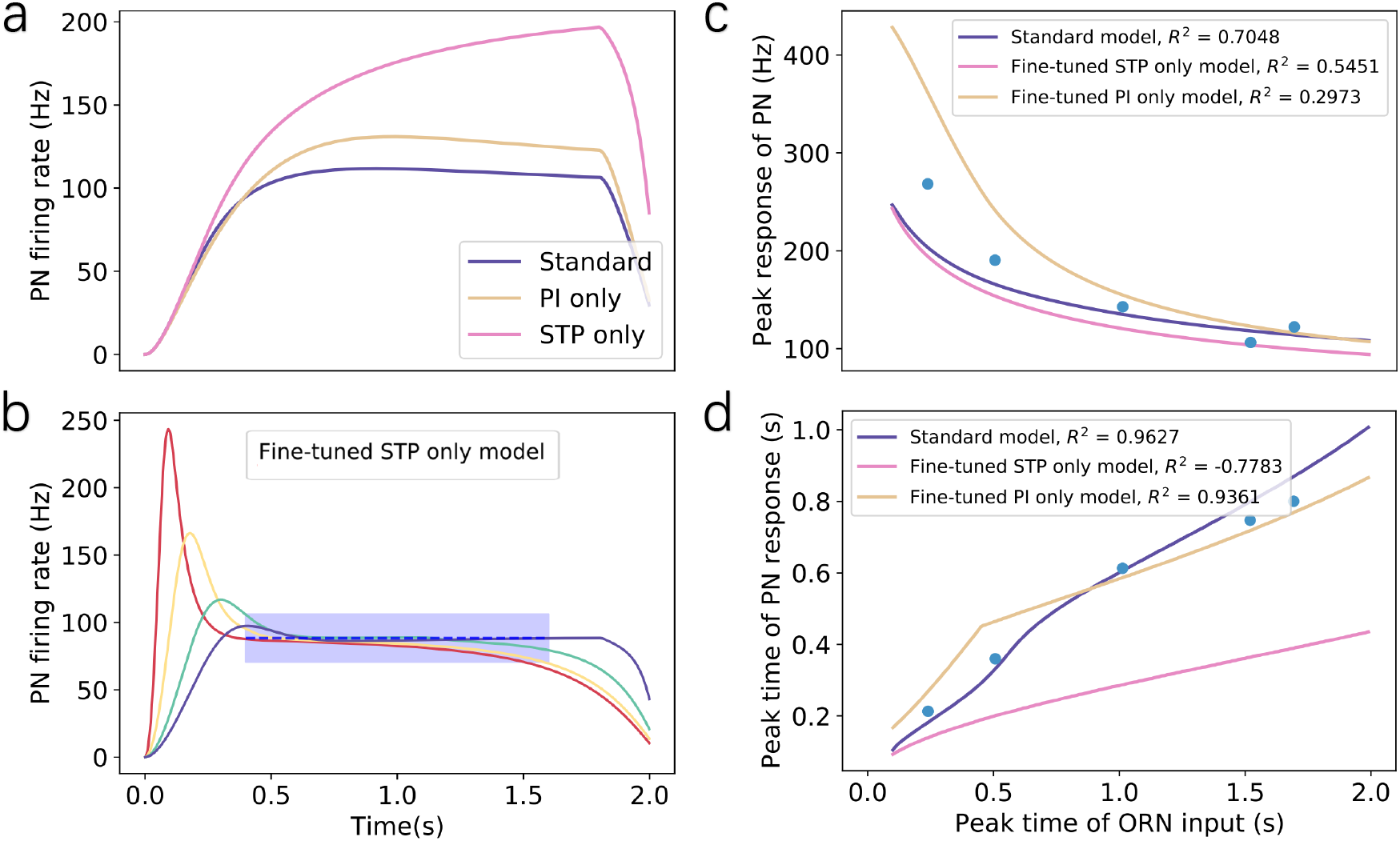
PN response under different conditions reveals the interaction between STP and presynaptic inhibition (PI). (a) PN response in three model variants to triangle-shaped ORN input as in Fig.5 with peak time at 1.8 s and peak amplitude 120 Hz. All the three models share the same set of parameters as in Fig. 5, except that the PI only model has *τ_D_* = *τ_F_* = 0 and the STP only model has *ρ* = 0. (b)The best fit to the experiments by using the STP only model. For all model variants, the parameters are selected to best fit the plateau height (shaded region) for the slowest input. Comparing the fitting of three model variants (standard, STP only, and PI only) with experimentally observed peak response (c) and peak response time (d), we found the standard model performs much better than other model variants. Note that the *R*_2_ value for the STP only model is negative, which means that the STP-only model fails to fit the peak time data when it is tuned to fit the plateau height as shown in (b). Parameters in the variant models that are different from the standard model are: *τ_D_* = 270 ms, *τ_F_* = 100 ms, *ρ* = 0, *U* = 0.2, *ω^EE^* = 90 nS for the STP-only model; *τ_D_* = 0 ms, *τ_F_* = 0 ms, *ρ* = 0.01, *τ_p_* = 250 ms, *U* = 0.25, *ω^EE^* = 80 nS for the PI-only model.

**Fig. S3.**
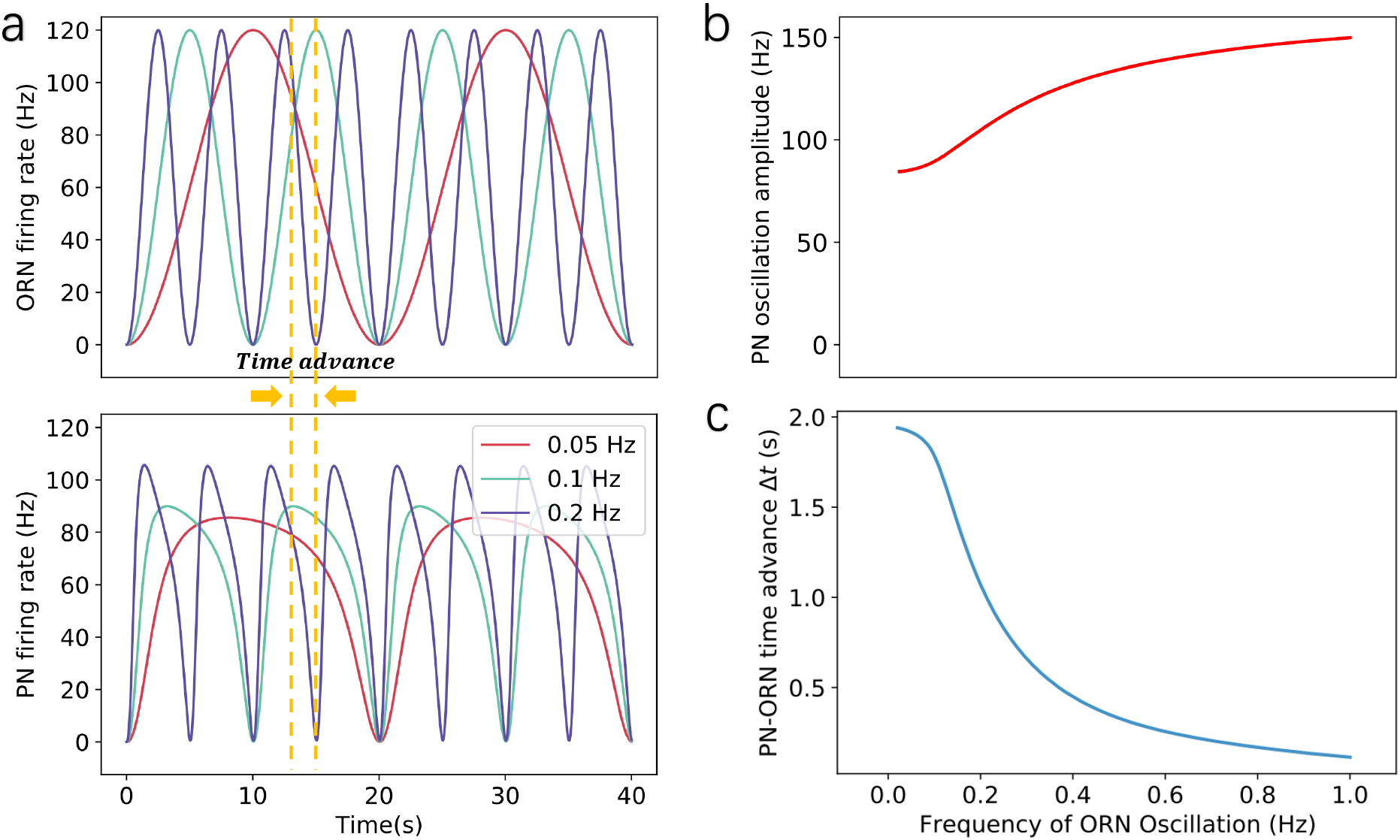
Prediction of PN response to oscillatory ORN inputs. (a) Upper panel: Simulated sine-wave ORNs firing rates with different frequency. The peak firing rates are the same for all inputs. Lower panel: responses of PNs to sine-wave ORNs inputs. The two dotted lines mark the peak of ORN input and its corresponding PN output with a frequency of 0.1 Hz. The gap between these dotted lines is defined as the time advance Δ*t* of PN’s response. (b) The peak responses of PNs increase with the frequency of ORN oscillation. (c) PN’s response always reaches to a peak earlier than that of the input signal, but the time advance Δ*t* decreases as the oscillation frequency increase. Parameters used in the model are the same as in Fig. 5

